# SelexGLM differentiates androgen and glucocorticoid receptor DNA-binding preference over an extended binding site

**DOI:** 10.1101/176073

**Authors:** Liyang Zhang, Gabriella D. Martini, H. Tomas Rube, Judith F. Kribelbauer, Chaitanya Rastogi, Vincent D. FitzPatrick, Jon C. Houtman, Harmen J. Bussemaker, Miles A. Pufall

## Abstract

The DNA-binding interfaces of the androgen (AR) and glucocorticoid (GR) receptors are virtually identical, yet these transcription factors share only about a third of their genomic binding sites and regulate similarly distinct sets of target genes. To address this paradox, we determined the intrinsic specificities of the AR and GR DNA binding domains using a refined version of *SELEX-seq*. We developed an algorithm, *SelexGLM*, that quantifies binding specificity over a large (31 bp) binding-site by iteratively fitting a feature-based generalized linear model to SELEX probe counts. This analysis revealed that the DNA binding preferences of AR and GR homodimers differ significantly, both within and outside the 15bp core binding site. The relative preference between the two factors can be tuned over a wide range by changing the DNA sequence, with AR more sensitive to sequence changes than GR. The specificity of AR extends to the regions flanking the core 15bp site, where isothermal calorimetry measurements reveal that affinity is augmented by enthalpy-driven readout of poly-A sequences associated with narrowed minor groove width. We conclude that the increased specificity of AR is correlated with more enthalpy-driven binding than GR. The binding models help explain differences in AR and GR genomic binding, and provide a biophysical rationale for how promiscuous binding by GR allows functional substitution for AR in some castration-resistant prostate cancers.

## INTRODUCTION

Gene expression programs are precisely regulated by transcription factors (TFs), a class of DNA-binding proteins that orchestrate the activity of the RNA polymerase II and chromatin-modifying complexes. The DNA binding domains (DBDs) of TFs fall into families consisting of dozens or even hundreds of members (Weirauch et al. 2014), leading to similar DNA sequence preferences among family members. Nonetheless, subtle quantitative differences in DNA binding specificity between related TFs are associated with large qualitative differences in the sets of targets genes they control (Maerkl and Quake 2007). To understand gene regulation and regulatory networks, it is therefore essential not only to accurately quantify these differences in DNA recognition but also determine the structural and physical basis of that specificity. The former can be done using comprehensive, unbiased experimental and computational methods; the latter requires more focused mechanistic analyses.

The intrinsic DNA binding specificities for hundreds of TFs have been profiled using a number of different high-throughput assays. These include (universal) protein binding microarrays (PBM)(Berger and Bulyk 2009), bacterial 1-hybrid (B1H) (Meng et al. 2005), and (solution-based) high-throughput SELEX (HT-SELEX) (Zhao et al. 2009; Jolma et al. 2010; 2013). These methods have been effective at defining core binding motifs (a sequence pattern that contains the main determinants of TF:DNA interaction), which can be represented as position weight matrices (Stormo 2000) or position-specific affinity matrices (Foat et al. 2006), and visualized as information content (Schneider et al. 1985) or energy/affinity logos (Foat et al. 2006). The models are available via databases such as Jaspar (Mathelier et al. 2015), UniProbe (Orenstein and Shamir 2014; Hume et al. 2015) and CisBP (Weirauch et al. 2014). Although for most TFs these approaches have been effective for defining a core sequence pattern required for high-affinity DNA binding, the resulting motifs often fail to discriminate between TFs from the same structural family.

Interactions with base pairs outside the core binding site can contribute to differences in DNA sequence preference between paralogous TFs (Zhou and O’Shea 2011; Maerkl and Quake 2007; Fisher and Goding 1992). For example, Cbf1p and Tye7p, members of the basic helix-loop-helix (bHLH) family in *S. cerevisiae*, recognize similar E-box core sequences (CANNTG), but distinct flanking preferences became apparent when binding was assayed over a larger footprint using custom-designed genomic context protein binding microarrays (gcPBMs) (Gordân et al. 2013). Differences in specificity associated with flanking regions have also been observed in HT-SELEX data (Jolma et al. 2013), and for ETS proteins using PBMs (Wei et al. 2010). Moreover, *SELEX-seq* technology has been used to show that Hox proteins read out the spacer sequences between half-sites in distinct ways when binding as heterodimers with the cofactor Exd (Slattery et al. 2011; Abe et al. 2015). However, although these techniques have demonstrated that flanking sequences are important, they cannot accurately quantify sequence specificity over larger footprints (> 15bp) without a specialized design (e.g. gcPBM). Thus, a more refined, general approach is needed to assess whether functional differences between TFs from the same family can be attributed to differences in intrinsic DNA binding specificity considered over the full TF:DNA interface.

The “missing specificity” issue – or the functional differences between transcription factor paralogs that cannot be explained by initial measurements of binding specificity – is brought into sharp relief by castration resistant prostate cancer (CRPC). Prostate cancer is driven by androgen signaling through regulation of gene expression by the androgen receptor (AR, *AR*) (Watson et al. 2015). Blocking of androgen synthesis and inhibition of ligand binding to AR have both been effective treatments. Unfortunately, CRPC eventually arises due to alternative production of androgens or activation of AR (Feldman and Feldman 2001). In some cases, CRPC is accompanied by increased expression of the glucocorticoid receptor (GR, *NR3C1*), which then functionally substitutes for AR by activating a subset of the AR transcriptional program that drives cancer progression (Arora et al. 2013). Despite this overlap, chromatin immunoprecipitation followed by sequencing (ChIP-seq) in LNCaP-1F5, a cell line model of CRPC, has shown that AR and GR share only about a third of their genomic binding sites (Sahu et al. 2013; 2014). Although co-factors such as the transcription factor FoxA1 help distinguish between AR and GR binding (Belikov et al. 2016; Sahu et al. 2013), these two factors still bind distinct loci in their absence suggesting an intrinsic ability to distinguish sequences that is not reflected in previous measurements of their in vitro specificities (He et al. 2012; Jin et al. 2014; Pihlajamaa et al. 2014; Jolma et al. 2013).

It is not clear how AR and GR, both members of the steroid hormone receptor (SHR) family, are directed to different genomic loci. Existing ChIP-seq and HT-SELEX (Nelson et al. 1999; Jolma et al. 2013; Sahu et al. 2013) studies suggest that AR and GR bind indistinguishable 15bp core motifs composed of inverted hexameric half-sites separated by a three-base-pair spacer: <u>RGAACA</u> NNN <u>TGTTCY</u>. Crystallographic evidence indicates that AR and GR each bind DNA as head-to-head dimers, with the two monomer subunits each occupying a half-site and dimerizing over the spacer (Shaffer et al. 2004; Meijsing et al. 2009; Watson et al. 2013). Our detailed analysis of AR- and GR-DNA crystal structures indicates that they share an identical DNA binding interface (**Fig. S1**). Conserved residues make specific contacts in the major groove at positions 2, 4, and 5 (Luisi et al. 1991; Arbuckle and Luisi 1995) (**Fig. S2**) accounting for most of the binding energy, although other non-contacted base pairs within the half-site have also been shown to affect GR affinity (La Baer and Yamamoto 1994). Additional energy is derived from backbone contacts along the 3bp spacer, which are sensitive to minor groove width in GR (Meijsing et al. 2009; Watson et al. 2013). Contacts made with DNA sequences flanking the core motif further contribute to GR affinity (Meijsing et al. 2009). In this work, we test whether, despite their conservation, AR and GR use this shared DNA-binding interface differently to distinguish sequences within and flanking the core 15bp motif.

Models that estimate the contribution of individual DNA features to the overall binding free energy by directly fitting PBM intensities have been shown to capture sequence specificity with greater accuracy than classification-based algorithms (Weirauch et al. 2013). The *MatrixREDUCE* algorithm (Foat et al. 2006) is an early example of this general approach that has been extended in other studies (Zhao and Stormo 2011; Gordân et al. 2013; Riley et al. 2015). Analysis of SELEX read counts, however, differs fundamentally from that of PBM intensities. and typically involves the tabulation of relative enrichment for all oligomers (“k-mers”) of a given length (Riley et al. 2014). Because the expected number of reads matching any particular k-mer decreases exponentially with *k*, generation of comprehensive tables is typically limited by sequencing cost to a 12-15bp footprint (Abe et al. 2015; Slattery et al. 2011; Jolma et al. 2013). Feature-based models have been used to interpret the oligomer tables, either in the form of a simple position weight matrix (Jolma et al. 2013) or based on a combination of base and shape features (Abe et al. 2015). Although this approach has provided valuable insights, it is limited by the restriction on binding-site size inherited from the oligomer tables on which they are based. Thus, in order to distinguish between the specificities of AR and GR over their full footprints, more refined experimental and computation techniques are required.

## RESULTS

### AR interacts with DNA over a larger footprint than GR

To determine the intrinsic specificity of AR and GR at high resolution, we performed *SELEX-seq* (Slattery et al. 2011; Riley et al. 2014) for homodimers of the DNA binding domain (DBD) of each factor (Fig. 1A, **Fig. S1C**). Although SELEX-seq (and lower throughput predecessors) have been developed over the years (Ogawa and Biggin 2011; Djordjevic 2010), increased sequencing and computational power have allowed some refinements. Purified proteins were incubated with a pool of DNA molecules, each containing a larger (23bp) random region than typical, flanked by Illumina adapters and tagged at one end with Cy5. Electrophoretic mobility shift assays (EMSA) were used to separate dimer-bound DNA sequences over eight rounds of affinity-based selection (Fig. 1A, **Fig. S3**). After each round, the concentration of the isolated DNA was quantified by qPCR, and then in parallel amplified for reselection and packaged into a sequencing library by adding Illumina flow cell adapters. Each library was then sequenced to a depth of ∼10^7^ reads. Preliminary analysis of the *SELEX-seq* data revealed unexpected differences between AR and GR. Following a previous study (Slattery et al. 2011), we estimated affinities as normalized oligomer enrichments, using the R/Bioconductor package *SELEX* (Riley et al. 2014). Biases in the initial round zero (R0) pool were estimated using a 5^th^-order Markov model, after which we computed the information gain between the initial (R0) and final round (R8) to estimate binding site size. For GR the information gain peaks at 15bp (Fig. 1C), consistent with the previously defined core motif. However, for AR information content continues to increase beyond 15bp (Fig. 1D), indicating a sensitivity to base identity over a larger binding site. One concern was that sequences would be over-selected after 8 rounds. Due to the diversity of the pool and lack of a strict consensus sequence for both AR and GR, very few 23-mers sequences are observed more than once in the sequenced libraries (Fig 1E). This indicates not only that the libraries were not over-selected, but also that further rounds of selection might allow better discrimination of lower affinity sequences without over-selecting high-affinity sites (Djordjevic 2010). A cursory k-mer based analysis indicated that AR and GR differ in their specificities, sharing only about a third of their significantly enriched 15-mers (Fig. 1F, **S4A,B**), with substantially different preferences among those common 15-mers (Fig. 1G), and confirmed specificity for AR over a larger footprint (**Fig. S4C**). Quantitative EMSAs supported the rank order of these sequences in terms of their enrichment in the *SELEX-seq* experiment (**Fig. S4E,F**).

**Figure 1:**
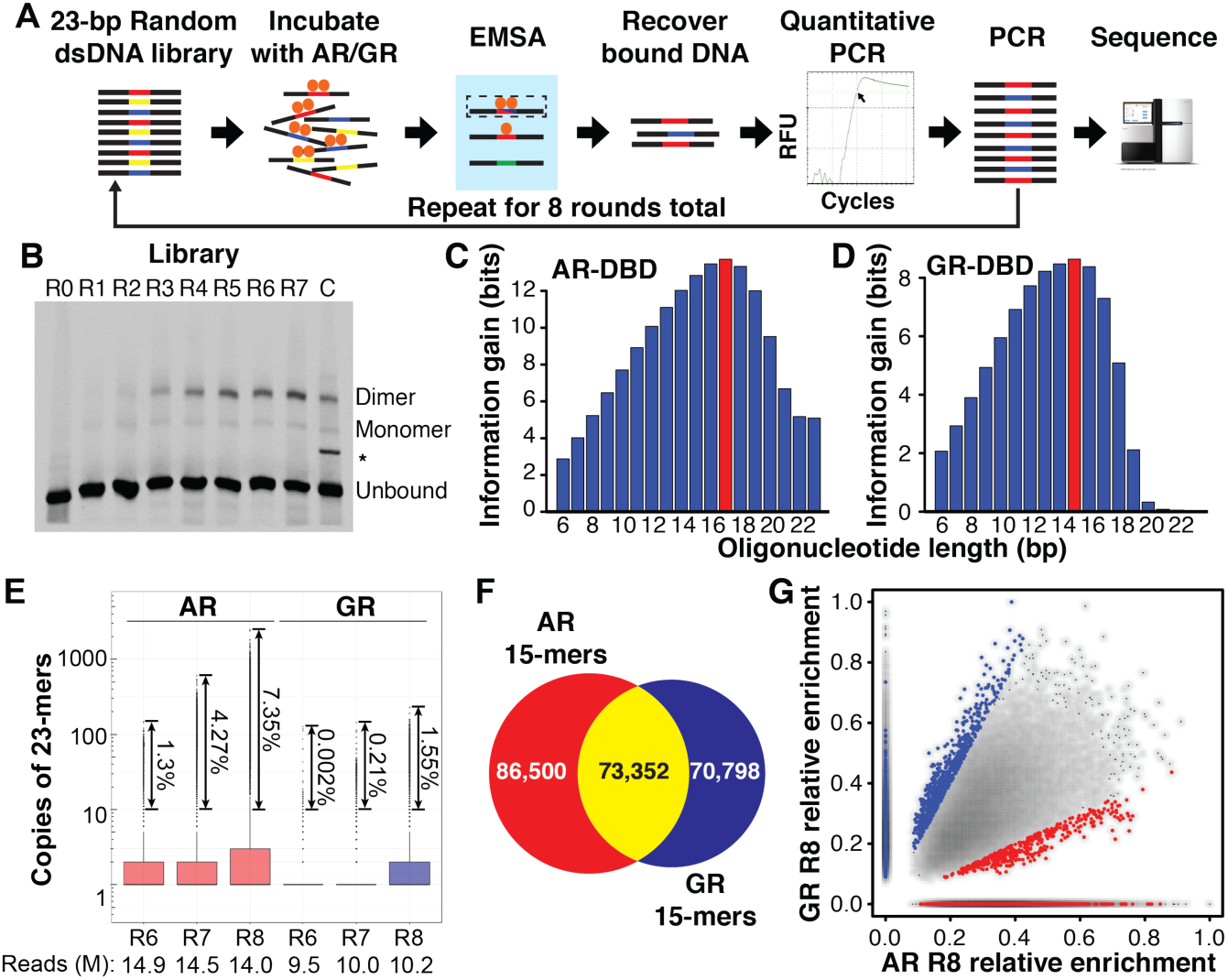
*SELEX-seq* reveals differences in AR- and GR-DBD DNA binding specificity. *SELEX-seq*. A 70bp ssDNA library with 23bp randomized region was incubated with the DNA binding domains (DBD) of AR or GR and separated into monomer and dimer species by EMSA. Dimer bound DNA was recovered, quantified by qPCR, amplified as the library for the next round, and repeated for eight rounds. Each round of library, including the initial dsDNA library, was sequenced. **(B)** EMSA gel showing the enrichment of dimer bound sequences after each round of selection for GR-DBD. The intensity of the shifted band plateaus after round 7. A high-affinity palindromic sequence served as a control to locate the dimer band. (*) marks an artifact during the synthesis of control sequence, but not observed in the SELEX library. **(C, D)** Information gain, or Kullback-Leibler divergence, from R0 to R8, as a function of oligonucleotide length. **(E)** Boxplot showing the multiplicity of unique 23-mers in each of the last three rounds of SELEX-seq selection for AR and GR. Even for the most highly selected library (AR R8) fewer than 10% of all reads have 10 copies or more, indicating that the libraries are not over-selected. (**F)** Venn diagram showing the overlap of sequences for AR- and GR-DBD with at least 100 sequencing counts. **(G)** Scatterplot of sequences that were commonly bound (yellow from panel **(F)**) by AR- and GR-DBD.

Further analysis indicated that both spacer sequence and flanking poly-A tracts contribute to AR affinity over at least 21bp (**Fig. S4C**). At that footprint size, it is not possible to accurately quantify differences in affinity using oligomer tables as in previous studies (Slattery et al. 2011; Jolma et al. 2013) for the following reasons. The most enriched 21-mer for AR (AAA<u>AGAACA</u>CGA<u>TGTACT</u>TTT) is contained in ∼4,000 reads out of the ∼10^7^ that make up the R8 library. Because the expected count for suboptimal 21-mers in R8 scales as the 8^th^ power of affinity, a mutated 21-mer sequence with an affinity 3-fold lower than the optimal 21-mer would have a 3^8^-fold (6,561) lower expected frequency in the pool, and therefore be unobservable. This is problematic, as much lower-affinity TF binding sites have recently been shown to be functionally important (Tanay 2006; Crocker et al. 2015; Farley et al. 2015). Furthermore, 15-mers covering only one of the two half-sites can be more strongly enriched than those centered at the dimer site for AR (see **Table S1** and caption of **Fig. S3F**), a misleading effect that arises because individual 15-mers cannot cover the entire binding site but their presence nevertheless can imply favorable flanking sequence. Although the logos suggest a larger footprint for AR, the k-mer tables (**Figure S4C,D**) do not allow for biophysical interpretation. Together, these limitations motivated us to develop a new computational technique to accurately describe DNA recognition over a large binding-site.

### Quantifying DNA binding specificity using feature-based modeling

To overcome the limitations of oligomer tables outlined above, we developed *SelexGLM*, a flexible modeling strategy based on Poisson regression that allows us to estimate binding free-energy contributions throughout the protein-DNA interface directly from the *SELEX-seq* read counts (Fig. 2). We use a standard equilibrium thermodynamics description of protein-DNA interaction that was previously used to analyze protein binding microarray (PBM) data (Foat et al. 2006; Zhao and Stormo 2011; Gordân et al. 2013; Riley et al. 2015). The relative binding free energy for sequence *S* is modeled as a sum of parameters ΔΔ*G*_*ϕ*_ associated with the DNA sequence features *ϕ* ∈ Φ(*S*) that characterize the difference between *S* and a reference sequence *S*_ref_ (typically the highest-affinity sequence):

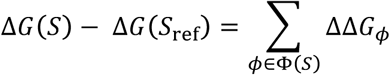

**Figure 2:**
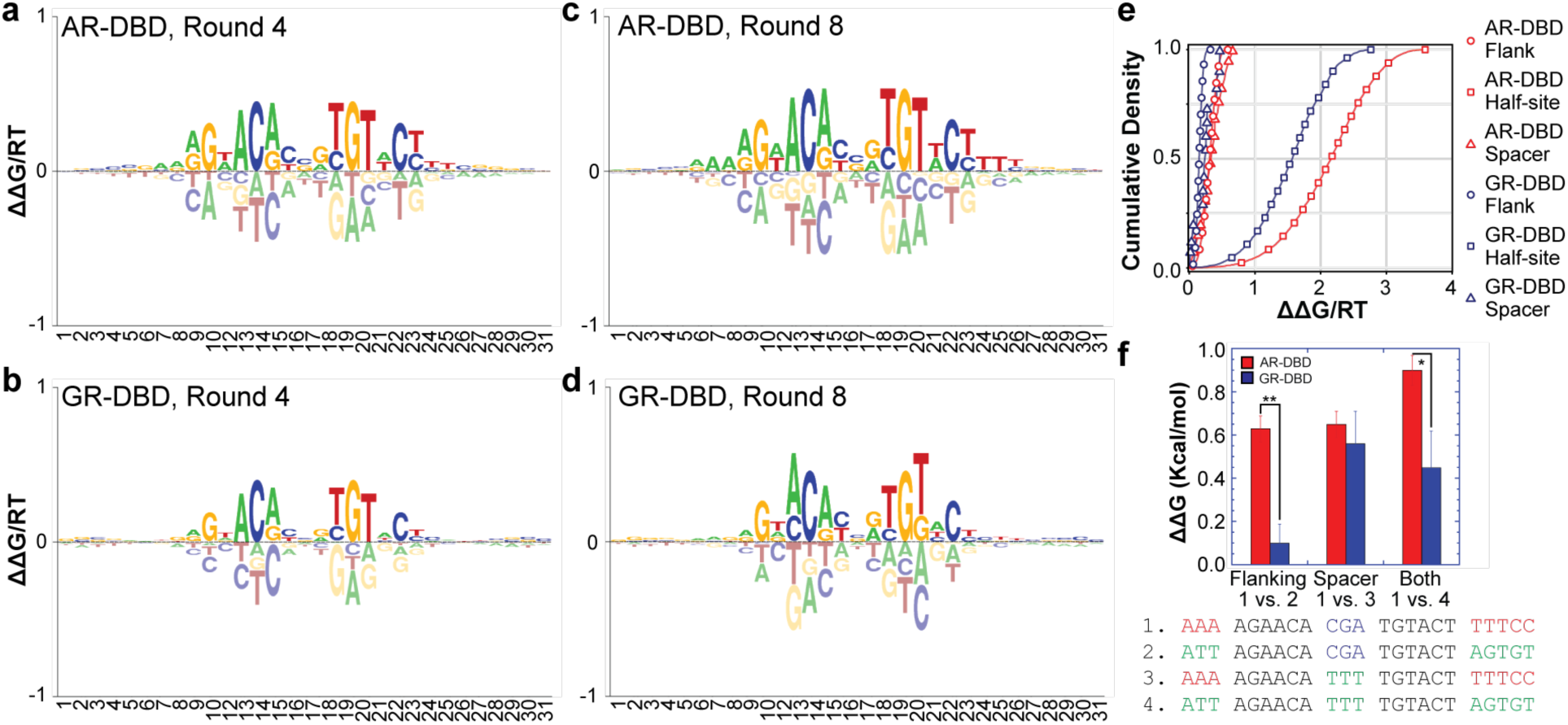
*SelexGLM* shows differences in DNA recognition between AR and GR throughout their binding sites. **(A-D)** Energy logos for AR-DBD (top) and GR-DBD (bottom), obtained by fitting biophysical models for protein-DNA interaction to the *SELEX* read counts using an iterative generalized linear modeling approach based on Poisson regression, implemented as *SelexGLM*. Highly similar logos were obtained using two separate rounds of data. See **Fig. S5** for logos generated using round 4 to 8 data. **(B)** Cumulative distribution functions for the contribution of half site (squares), spacer (triangles), and flanking (circles) sequences on AR-DBD (red) and GR-DBD (blue) binding energy. **(C)** Validation of the contribution of flanking A tracts and spacer to AR- and GR-DBD binding performed by quantitative electrophoretic mobility shift assay. Loss of flanking A tracts is more detrimental to AR-than GR-DBD (1 vs. 2), whereas changing spacer can have detrimental effects on the binding of both (1 vs. 3). Error bars are the standard error of the mean based on at least 3 repeats of each experiment. (* P-Value ≤ 0.05, ** P-Value ≤ 0.01, t-test)

In this study, we restricted ourselves to single-nucleotide features (e.g., *ϕ* = *C*_3_, denoting the presence of a C at position 3 within the binding site window). Our modeling assumptions imply that the combined effect of multiple mutations within the binding site is additive in terms of binding free energy or multiplicative in terms of relative affinity.

The model parameters are found through an iterative fitting process that starts from an initial estimate, or seed. To generate this seed, we construct a relative enrichment table using a footprint large enough to capture the binding site core but not so large that counts get too low. We chose a seed length of 15bp for both AR and GR, but the final model is only weakly dependent on this choice and can have a much larger footprint (31bp in our case; see below).

The negative logarithm of the relative enrichment of each mutated 15-mers is used as an initial estimate of ΔΔ*G*_*ϕ*_/*RT* for each feature.

Once seeded, the model is refined by alternating between two steps. In the first step, we determine the highest-affinity binding site within each unique observed SELEX probe in the data (“affinity-based alignment”). This allows us to construct a design matrix *X* defining each DNA feature (in this case each base pair) relative to the optimal binding window in each probe; only probes whose rate of selection is dominated by a single binding site offset are included (see **METHODS**). In the second step, the design matrix is used to fit a generalized linear model (GLM) to the read counts, leading to a re-estimated set of free-energy coefficients:

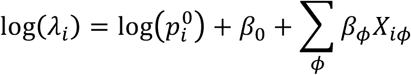

Here λ*_i_* is the expected value of the read count *y* for probe *i*:

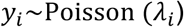

Each model coefficient β_*ϕ*_is used as a re-estimate of ΔΔ*G*_*ϕ*_/*RT*, which are then used to update the design matrix, and the process repeats. Convergence is reached once the position of the optimal binding window no longer changes for any of the probes in the data set after re-estimation of the free-energy coefficients.

### Earlier rounds of selection are sufficient for SelexGLM analysis

We performed SELEX-seq over eight rounds of selection in order to obtain linear enrichment of sequences appropriate for k-mer enrichment based analysis. When we analyzed each round of selection through SelexGLM, we were surprised to find that, although R8 provided the highest-resolution model without over selection (Fig. 1E, **Fig. S5A-J)**, high-quality models could also be generated from earlier rounds of selection **(Fig. S5A-J)**. However, the accuracy of the models increases in the later rounds, particularly when it comes to distinguishing the base-pair substitutions with the most deleterious effect on binding **(Fig. S5, S6)**. Thus we used models generated from R8 for both proteins in all subsequent analyses.

### AR is distinguished from GR by a preference for poly-A sequences outside the 15bp core

We used *SelexGLM* to build recognition models as position specific affinity matrices (PSAMs) for AR and GR with a 31bp footprint, significantly larger than the 15bp binding site core and even the 23bp variable region of the SELEX probes. *SelexGLM* is capable of fitting such wide models because it considers offsets within the probe that partially cover the fixed sequences upstream and downstream of the variable region. The corresponding energy logos for these PSAMs confirm the previously modeled 15bp size of the GR binding site (Fig. 2B,D), but reveal AR’s preference for poly-A sequences flanking the 15bp core (Fig. 2A,C) and surprising differences within the 3bp central spacer (Fig. 2E, and more evident in **Fig S4C,G,H**). For AR, replacing the best flanking sequence (AAAA) with the worst (TCGC) on one side leads to a 1.6-fold reduction in affinity (or 2.6-fold when both flanks are replaced) (Fig. 2E). Validation by qEMSA confirmed that spacer sequence affects the affinity of both AR and GR, but that flanking sequences affect only AR (Fig. 2F). The observed change for AR (ΔΔG = 0.60 ± 0.05 kcal/mol) was larger than predicted (ΔΔG = 0.26 kcal/mol), perhaps because our model ignores dependencies between nucleotide positions within the binding site.

### Further differences in AR and GR specificity are encoded throughout the 15bp core

Though the AR and GR binding models are similar at first glance, detailed analysis of the PSAMs highlights AR’s sensitivity to changes from the consensus sequence in the central 15bp core region (Fig. 3a). Based on these differences, we tested sequences that favor AR (Fig. 3B, **Table S2**, Shape-1 and Shape-2) and GR binding (Fig. 3B, **Table S2**, Shape-4) and verified that the two proteins can differentiate between DNA sequences over an order of magnitude in affinity (Fig. 3B, **Figure S5D**). As the two proteins have identical base-reading chemistry at the DNA-binding interface, we examined whether their sequence preferences could be explained in terms of DNA shape readout. Indeed, the shape preferences of AR and GR contrast sharply: the AR benefits from a narrowed minor groove in the flanking regions (Fig. 3C) whereas the GR prefers binding to sites with widened minor grooves within the half-sites (Fig. 3D). To validate this finding, we tested a sequence with a half-site predicted to have a narrow minor groove (TTTTAT) (Zhou et al. 2013) and found that GR bound significantly worse than AR (Fig. 3B, **S5D**, Shape-3). Similarly, we tested a site predicted to have a wide minor groove (GGGACA, Fig. 3B, **S5D**, Shape-4) and found a preference for GR. More dramatic examples of sequences predicted to have disparate GR and AR affinities are plentiful in the low affinity range (K_D_ ≥ 5μM), however measurements in this range are near the non-specific binding limit and have not been reliable (data not shown). In addition to differences in core preference, the nearly inverted minor groove preferences in the flanks for AR and the half-site for GR suggest the two proteins have different recognition modes.

**Figure 3:**
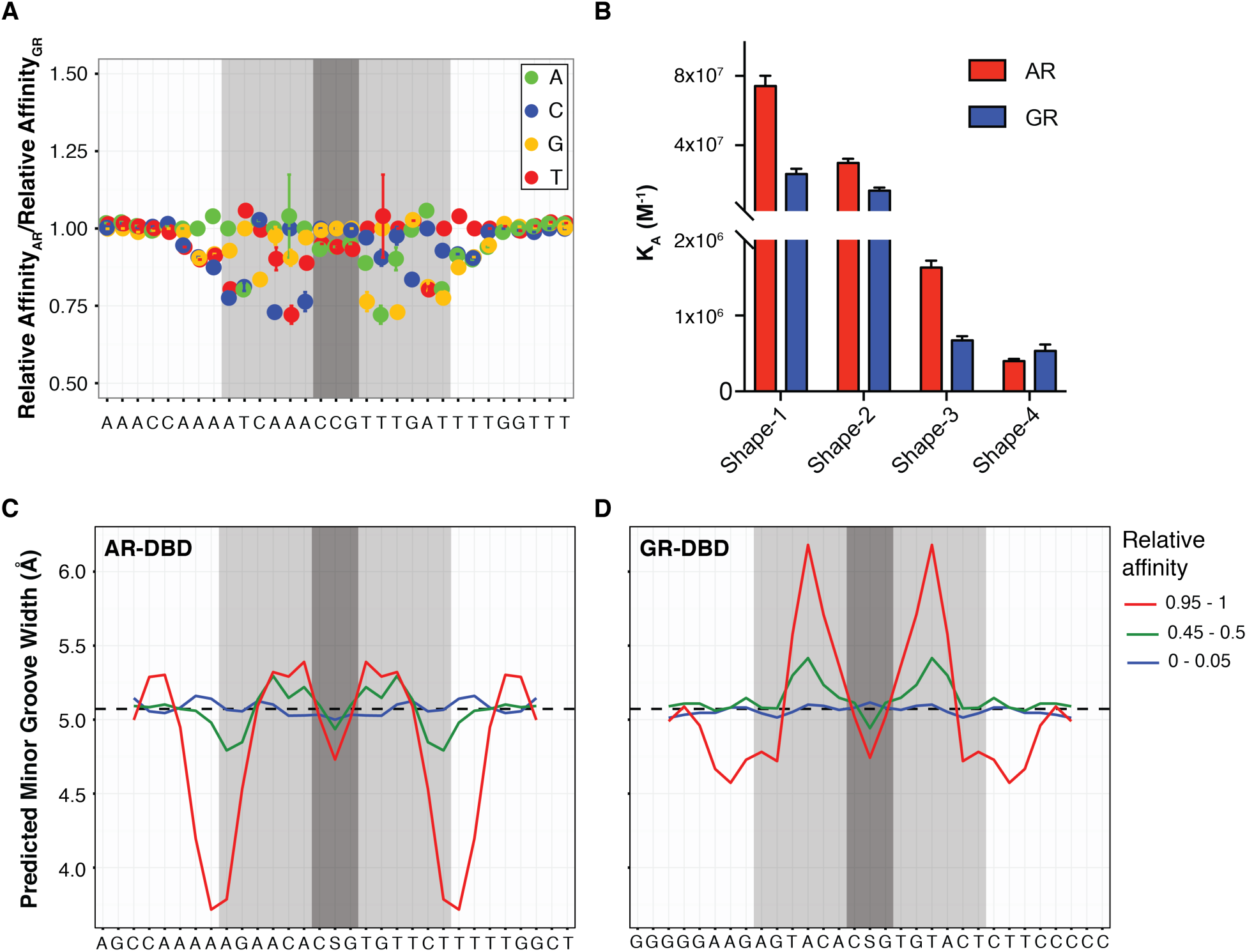
Difference in DNA shape readout between AR and GR. **(A)** Difference in ΔΔG/RT values between AR and GR at each nucleotide position, normalized by their mean across all four bases. **(B)** Quantitative EMSA was used to measure the affinities of AR- and GR-DBDs for four sequences that maximally favor AR (Shape-1) and GR (Shape-4) and test the importance of half-site minor groove width (Shape-3 and −4). Each measurement was performed at least three times. Error bars represent SEM. **(C)** Average minor groove width profile for the top /middle/bottom 5% of sequences in terms of affinity for AR and **(D)** for GR.

### The thermodynamics of binding reflect the specificity of AR and GR

To understand the thermodynamic basis of the differing AR and GR binding modes, we measured the effect of varying flanking and spacer sequences using isothermal titration calorimetry (ITC). Our ITC data allowed us to accurately quantify the affinity (and thus ΔG) of bound sequences and the contribution of individual sequence features to the enthalpy (ΔH) and entropy (ΔS) of binding (Buurma and Haq 2007). Although the *K*_*D*_*s* for AR and GR sequences were consistent with the PSAMs derived from the SELEX data (**Fig. S7C**), including AR preference for poly-A flanks, it was immediately apparent from the heat of binding (Fig. 4A) that AR and GR recognize the same DNA sequences differently. Overall, for the sequences tested, AR binding is more enthalpically driven (low ΔH) and GR binding more entropically driven (high TΔS) (Fig. 4C,D). This is consistent with previous findings; TF families engaged in direct readout via the major groove are more enthalpically driven whereas those engaged in readout via the minor groove or backbone contacts are more entropically driven by solvent exclusion (Privalov et al. 2011; 2007). Our data also identify an exception to this general pattern: AR’s recognition of the poly-A stretches in the minor groove is driven not entropically, but rather enthalpically. This observation contributes to our understanding of specificity: enthalpic contributions to binding are not solely the result of hydrogen bonds with individual base-pairs or backbone phosphates, but can also result from interaction with a DNA feature; the narrowed minor groove (Fig. 3).

**Figure 4:**
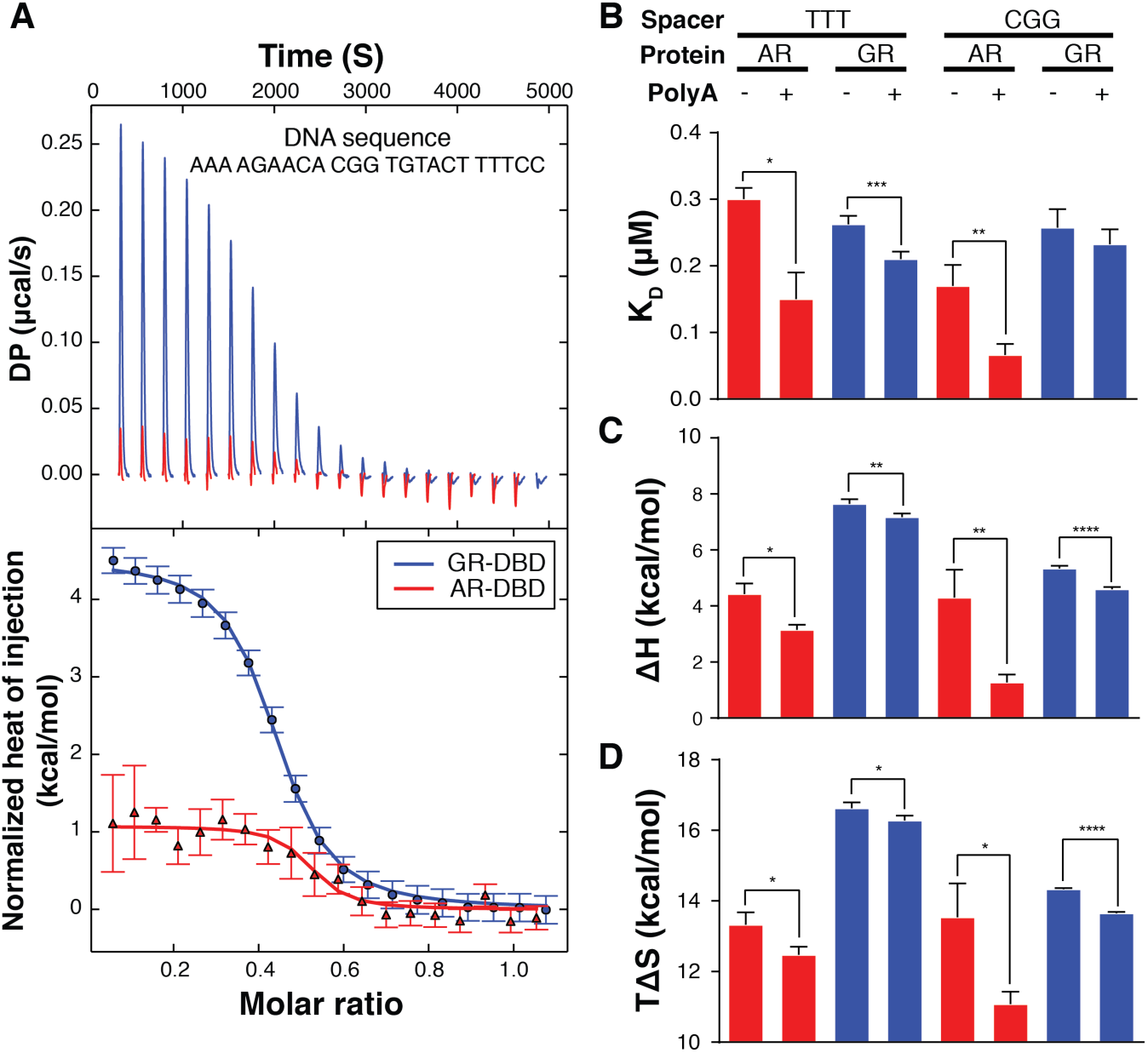
ITC analysis reveals distinct DNA binding thermodynamics between AR- and GR-DBD. **(A)** The raw heat titration signals (top) and normalized (NITPIC) heat of injection profiles (bottom) of AR- and GR-DBD bound to a given DNA sequence. Standard errors are estimated by NITPIC. **(B)** The K_D_ of AR-DBD and GR-DBD for four sets of sequences fit from the ITC data. AR-DBD affinity is increased with flanking As and an optimal spacer, whereas GR is insensitive. **(C)** Enthalpy, ΔH, is calculated from the heat of binding for each DNA sequence. Flanking sequences decrease ΔH for AR-DBD, enhancing affinity. Smaller indicates a greater contribution to affinity. **(D)** Entropy, ΔS, is calculated from the K_D_ (thus ΔG) and ΔH. GR-DBD affinity is more entropically driven. Larger is indicates a greater contribution to affinity. (* P-Value ≤ 0.05, ** P-Value ≤ 0.01, *** P-Value ≤ 0.001, t-test) Error bars represent the standard deviation from at least 3 experiments.

### Promiscuous GR specificity predicts binding at AR genomic loci

We next asked whether the *SelexGLM* models can explain why GR is able to functionally substitute for AR despite having non-overlapping binding sites in models of CPRC. To this end, we analyzed ChIP-seq data from LNCaP-1F5, a cell line model of prostate cancer engineered to overexpress GR and enable genomic mapping of both AR and GR binding under similar cellular conditions (Sahu et al. 2013; 2014). As a positive control, we confirmed that the *SelexGLM* models can differentiate ChIP-seq peaks from adjacent regions (area under ROC-curve 0.78 and 0.83 for AR and GR, respectively; Fig. 5A). Further, we observed a significant quantitative relationship between predicted affinity and degree of genomic occupancy (Fig. 5B), with ChIP-seq peaks in the top decile for *in vitro* affinity being significantly higher than those in the bottom decile (1.4-fold, p=7.4*10^-49^, Wilcoxon rank-sum test, for AR; 1.6-fold, p=2.8*10^-68^, for GR). To test whether the *difference* in intrinsic binding specificity between AR and GR discovered using *SELEX-seq* was reflected in the genomic occupancy patterns probed using ChIP-seq, we let our AR and GR PSAMs compete in a multiple linear regression model. As expected, when analyzing the variation in GR peak height in this manner, we found that the regression coefficient for GR affinity was significantly larger than that for AR (p<10^-6^, t-test; Fig. 5C) and that the latter did not deviate significantly from zero (p=0.97). However, when analyzing the variation in AR peak height, both the AR and GR coefficients were non-zero (p<10^-5^ and p<10^-7^, respectively) and did not significantly differ from each other (p=0.16), indicating that both AR and GR are likely to bind well at AR loci.

**Figure 5:**
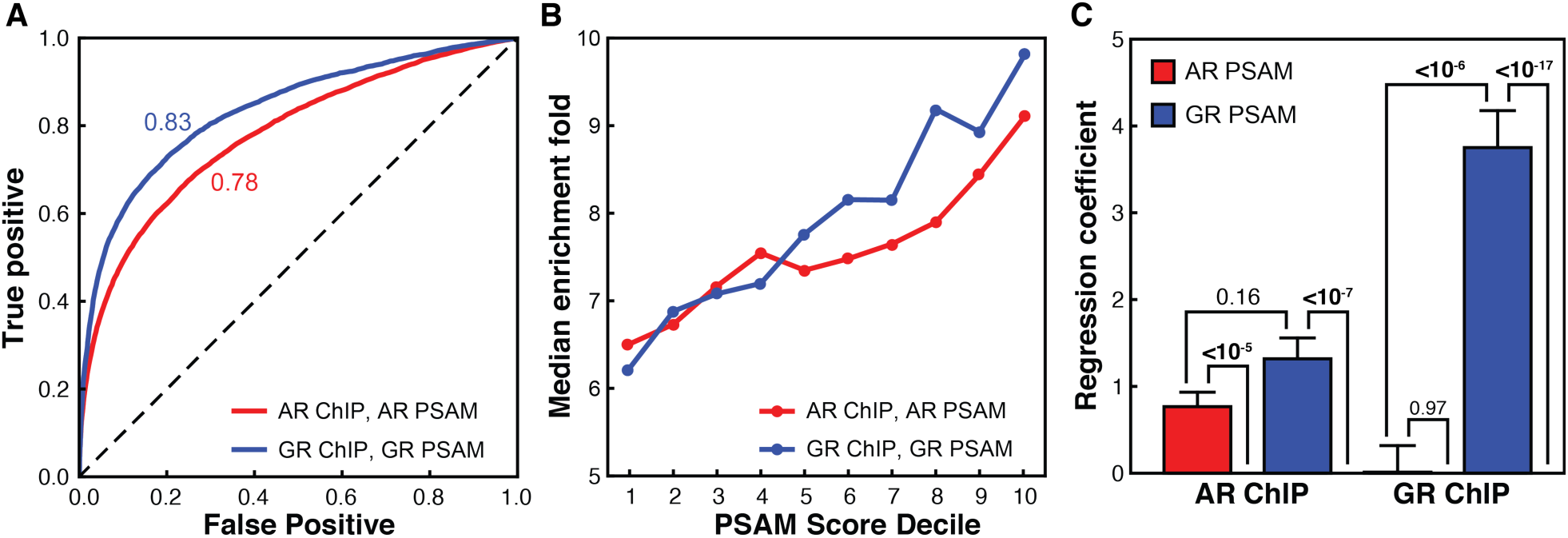
Differences of intrinsic specificity between AR- and GR-DBD are reflected in the respective cellular genomic binding profiles. **(A)** Ability of the AR (red) and GR (blue) PSAMs to identify true ChIP-seq peaks as measured by the receiver operator characteristic (ROC) of binary classifiers identifying peaks vs. adjacent regions. **(B)** Comparison of the median enrichment over background from ChIP-seq (y-axis) to the relative affinity for the strongest sequences within the ChIP-seq peak calculated from the position specific affinity matrix for AR (red) and GR (blue). In vitro affinity is binned by decile (e.g. 10 is the sequence space representing the top 10% highest affinity sites) **(C)** Multiple linear regression coefficients in models that use AR (red) and GR (blue) PSAM scores to predict AR (left) and GR (right) ChIP-seq peak enrichments. *p*-values were calculated using *t*-tests.

## DISCUSSION

In this study, we developed and validated a strategy for quantifying the DNA binding specificity of TFs with high precision over a much larger footprint than was previously feasible for SELEX data. This required experimental refinements to *SELEX-seq* and a new computational method, *SelexGLM*, to fit the free-energy parameters of a biophysically motivated model to the *SELEX-seq* data. Experimentally, we designed a library to accommodate the footprint of AR and GR defined by crystal structures (Meijsing et al. 2009; Shaffer et al. 2004), isolated dimer-bound DNA from a mixed population by EMSA, and performed qPCR between rounds of enrichment to avoid PCR artifacts that add noise to measurements. These refinements revealed that, despite their similarity, AR and GR have different sequence specificity both within and outside the core 15bp sequence that can be genetically tuned over a significant range, even in the absence of any co-factors. Compared to HT-SELEX assays performed on similar protein fragments, both of these refinements improved the accuracy of the model **(Fig. S8)**. SelexGLM was able to distinguish specificity over a 23bp footprint from this data **(Fig. S8A-D)**, whereas k-mer-based analysis only revealed a partial binding site at lower resolution **(Fig. S8E-G)**. Importantly, the models derived from HT-SELEX correlated poorly with the EMSA validation **(Fig S8H-J)**, and were less consistent and accurate than those derived from our SELEX-seq data.

The intrinsic specificity, thermodynamics of binding, and genomic localization indicate that GR is more promiscuous than AR, allowing it to accommodate a wider range of sequences. These binding properties are consistent with the behavior of GR in some CRPCs. Anti-androgen therapy can result in de-repressed GR expression, increasing the amount of GR in the cell (Arora et al. 2013; Watson et al. 2015), which, in the presence of endogenous or administered glucocorticoids results in an increase in the active concentration of GR in the nucleus. Our data is consistent with the idea that the more relaxed specificity provided by entropy-driven binding allows the excess GR to bind related sequences with reasonable affinity, including those previously bound by AR (Fig. 5C), enabling regulation of AR-driven genes aberrantly by GR (Arora et al. 2013). Thus, the biophysical properties of GR provide a rationale for how it can effectively substitute for AR in the context of CPRC.

The differences between AR and GR that our analysis uncovered were surprising, since the DNA binding interfaces of AR and GR are chemically identical. However, the integration of SELEX analysis and thermodynamic quantification of binding energies using ITC make it clear that AR and GR use these interfaces differently. Having comprehensively measured the specificity of AR and GR using all possible DNA ligands, it is clear that GR derives more entropic energy for the same DNA than AR. Since the majority of entropic energy is derived from solvent and ion exclusion from an interface, and the protein and DNA surfaces at the interface are chemically identical, we expected this contribution to be similar for AR and GR. However, the differences between GR and AR suggest an increase in conformational entropy within GR (1994; Frederick et al. 2007) (Fig. 4D), which would allow it to accommodate more diverse DNA sequences and shapes, but decrease its specificity. AR, conversely, does not derive as much binding energy from entropy, suggesting that precise positioning of hydrogen bonding in the half sites is more important. AR also derives enthalpic energy from the recognition of narrowed minor grooves created by poly-A flanking sequences. Our results provide direct evidence that the increased negative electrostatic focusing (Rohs et al. 2009) associated with a narrowed minor groove (Yoon et al. 1988) in the presence of interactions with basic residues (**Fig. S1, S2**) drives an increase in affinity. These differences in the thermodynamics of recognition must be mediated by non-conserved amino acids within the fold of the protein, indicating a difference in how the interface is scaffolded. This phenomenon has also been observed in recent studies of the C2H2 Zinc finger family of TFs (Hughes 2011; Persikov et al. 2015), and indicates that both the residues displayed and how they are configured are critical to distinguishing how transcription factor family members recognize different DNA sequences. It also suggests that, much as in the case of RUNX1 (Yan et al. 2004) and Ets-1 (Pufall et al. 2005), the conformational entropy of GR may provide an opportunity for regulation that could direct it away from AR sites.

To enable analysis of the *SELEX-seq* data over a large enough footprint, it was necessary develop an improved algorithm for analyzing *SELEX-seq* data, which we named *SelexGLM*. In a self-consistent procedure that echoes Gibbs sampling for discriminative motif discovery (Lawrence et al. 1993), *SelexGLM* alternates between an “affinity based alignment” step that identifies the dominant binding site within each sequenced DNA ligand, and a GLM fitting step that estimates the binding free-energy change associated with each possible base substitution in the DNA binding site, until convergence is reached. The Poisson statistics that underlie the GLM fits allow us to obtain precise estimates of the free-energy parameters even when individual unique DNA have very few counts, since each possible base substitution or “feature” can occur in many different contexts. Using a modest sequencing depth of ∼10^7^ reads per round, we were able to achieve precise, validated models over a footprint of 31bp that would accommodate many other transcription factors complexes.

*SelexGLM* also opens new avenues for studying the modulation of transcription factor function. Transcription factor binding and activity are regulated by post-translational modifications (Pufall et al. 2005), cofactor binding (Chodankar et al. 2014), and combinatorial control through interaction other transcription factors on DNA. The increased footprint size made possible by our method enables measurement of specificity for larger, heterologous complexes on DNA, and the effect of sequence on their occupancy. Further, *SelexGLM* allows us to quantify relatively modest effects on specificity distributed nature over the entire footprint, which evolution may exploit to fine-tune expression levels in a flexible manner, and which may be altered by phosphorylation of the TF or cofactor binding (Kumar and Calhoun 2008; Chodankar et al. 2014). Thus, our approach may enable us to begin to understand to what extent signal-dependent changes in expression are due to altered transcription factor specificity.

## METHODS

### Expression and purification of GR/AR-DBD

DNA sequences encoding the DNA binding domains (DBDs) of human Glucocorticoid receptor (GRα: 418-506) and Androgen receptor (AR-B: 557-647) were cloned into an N-terminal his_6_-tagged vector (pET28a, Novagen). Vectors were transformed into BL21DE3 Gold *E.coli* (Agilent) cells, and grown to an OD600 of between 0.2 and 0.4. The temperature was reduced to 27°C and 10 μM of ZnCl_2_ was added to the culture. Expression of recombinant protein was induced with 0.5 mM IPTG for 4 hours when OD600 reaches 0.6 to 1. Cells were then spun down at 6000g for 15 minutes, then resuspended in Ni^2+^ column loading buffer (25 mM TrisHCl, pH 7.5, 500 mM NaCl, 15 mM Imidazole, 1 mM DTT, 1 mM PMSF), snap frozen, and stored at −80°C until purification. Cell suspensions were lysed with an EmulsiFlex C3 homogenizer running at 15,000 psi. After three passes or more through the EmilsiFlex, soluble protein was isolated by ultracentrifugation at 40,000 rpm using Beckman Ti-75 rotor for 1 hour at 4°C then collecting the supernatant. The supernatant was loaded onto nickel affinity column (GE Healthcare Life Sciences) pre-equilibrated with 25 mM TrisHCl, pH 7.5, 500 mM NaCl, 15 mM Imidazole. Unbound protein was washed off the column with equilibration buffer, followed by a low imidazole bump (25 mM TrisHCl, pH 7.5, 500 mM NaCl, 30 mM Imidazole) to remove non-specifically bound protein. A linear gradient of imidazole from 30 mM to 350 mM was used to elute DBD. Fractions containing DBDs were pooled and dialyzed overnight at 4°C in 20 mM TrisHCl, pH7.5, 50 mM NaCl, 2.5 mM CaCl_2_ and 1 mM DTT. Thrombin was used to cleave the his_6_-tag during the dialysis. Dialyzed protein was ultracentrifuged (40,000 rpm, 1 hour, 4°C), and loaded onto a cation exchange column (HiTrap SP HP, GE Healthcare Life Sciences) pre-eliqulibrated with 20 mM TrisHCl, pH 7.5, 50 mM NaCl and 1 mM DTT. The DBD was eluted in a linear gradient of NaCl from 50 to 350 mM over 20 CVs. Fractions containing DBD were pooled, concentrated (Amicon Ultra – 3K, Millipore), filtered (Ultrafree®-CL), and monomers isolated by gel filtration (16/600 Superdex 200 PG, GE LifeSciences) in 20 mM HEPES, pH 7.7, 100 mM NaCl, 1 mM DTT. DBDs were collected and dialyzed against storage buffer (20 mM HEPES, pH 7.7, 100 mM NaCl, 1 mM DTT, 50% Glycerol), quantified (280 nm, ε = 5095M^-1^cm^-1^ for both AR-and GR-DBDs).

### SELEX Library design and synthesis

We designed our SELEX library to contain a 23bp random region flanked by primer binding regions conforming to the Illumina TruSeq Small RNA format for a total of 70bp. The library was ordered in 1 mmole format from IDT as single strand with handmix option over the randomized region. The complementary strand was synthesized by Klenow extention using 5’-Cy5 labeled TSSR1 primer (**Table S2**) on a PCR machine. Briefly, a reaction containing 2.5 μM ssDNA library, 5 μM 5’-Cy5-TSSR1, 150 μM dNTPs in NEB buffer 2 was incubated at 94°C for 3 minutes, and then gradually cooled down to 37°C over 45 minutes. Six units of Klenow were added to every 25μl of reaction, incubated at 37°C for 60 minutes, 72°C for 20 minutes, and gradually cooled down to 10°C over 45 minutes. dsDNA was purified using MinElute PCR column (Qiagen) and quantified by absorbance at 260 nm.

### SELEX-seq

#### Selection

To ensure at least 1X coverage of all possible 23mers, SELEX was carried out in a 120 μl binding reaction containing 0.63 μM purified DBD and 1 μM DNA library. At this size, there are ∼ 7.2×10^13^ DNA molecules in the reaction, representing ∼1.02X coverage of all possible 23mers (7.03×10^13^). Binding was carried out in a buffer that approximates the salt and crowding of the nucleus for 1 hour at 4°C (20 mM TrisHCl, pH 8.0, 150 mM KCl, 5% Glycerol, 1 mM EDTA, 5 mM MgCl_2_, 40 ng/μl PolydIdC (Sigma: P4925), 200 ng/μl BSA, 1 mM DTT, 200 mg/ml Ficoll PM400 (Sigma: F4375)). The reaction was then run out in multiple wells of a 10% native polyacrylamide gel (19:1 Acrylamide/Bis-acrylamide) in 0.5X TB containing 150μM MgCl_2_ (89 mM Tris-Boric acid, 150 μM MgCl_2_, pH 8.3) at 4°C. In order to ensure that SHR-DBD:DNA complexes were trapped, the sample was loaded while running at 200V to minimize the dissociation before entering the gel. DBD:DNA complexes were visualized using Cy5 fluorescence (GE ImageQuant LAS4010), isolated by excision, and bound DNA isolated by electroelution (Novagen D-tube dialyzer, 3.5 kDa) into native PAGE running buffer described above. Resulting DNA sequences were then purified using (Qiagen MinElute PCR clean-up), eluted (10 mM TrisHCl, pH 8.0) to a final volume of 180μl, and amplified to generate next round of library as described below. Because of the relatively low affinity of the SHRs for DNA, we were unable to shift enough DNA using a limiting amount of protein (<1:5) to allow PCR generation of pools for subsequent rounds without generating high molecular weight artifacts. We therefore incubated the library with a high protein:DNA ratio (0.63 μM:1 μM) to select all potential binding sequences in early rounds. Please note that this protein:DNA ratio requires more rounds of selection to begin linear enrichment of sequences. Since submission of this manuscript we have begun using a 1:10 protein:DNA ratio. It is critical to perform a size-selection (8% 1X TG gel) from the recovered bound DNA to remove the high molecular weight DNA that co-migrates with complexes prior to re-amplification. Together with controlled amplification cycle by qPCR (see below), this improved SELEX protocol is artifact-free, saves a few rounds of selection and is less saturated at early rounds. Typically, 4∼5 rounds of selection are sufficient for factors with long binding footprints (∼ 25-30 bp).

#### Library regeneration

To generate enough DNA (∼10^13^ molecules) to perform each round of SELEX and sequencing library prep, the recovered DNA had to be carefully amplified. As these libraries were susceptible to amplification artifacts caused by over-amplification (different type of artifact than those co-migrating with the complex), the optimal number of PCR cycles was first determined by qPCR. Briefly, 1 μl of recovered dsDNA was analyzed in 50 μl qPCR reaction (0.5 μM TSSR0 primer (**Table S2**), 0.5 μM unlabeled TSSR1 primer, 200 μM dNTPs, 0.1X SybrGreen (Invitrogen: S-7563), 0.5 unit of Phusion polymerase in 1X NEB Phusion HF buffer). The amplification curve was then analyzed to determine the maximal number of rounds within the linear amplification range (Typically less than 16 PCR cycles, depending on the amount of the template). Subsequently, 170 μl recovered library was divided into 85 reactions at 100 μl, and amplified with the determined cycle numbers (with 5’-Cy5-TSSR1). The amplification reactions were then combined, purified, and concentrated (Qiagen MinElute), and eluted with 45μl EB. The resulting libraries were then quantified and mixed into a new binding reaction as described above. Additional agarose gel electrophoresis is required to remove polydIdC if it is present in the recovered dsDNA, as the polydIdC interfered with PCR.

#### Library sequencing

To sequence the resulting libraries on the Illumina HiSeq platform, additional adapter sequences were added by limited-cycle PCR. Briefly, 400 ng of each SELEX library was amplified using the 5’ adaptor primer (TSSR2, **Table S2**) and the 3’ adaptor and barcoding primer (TSSR-RPIX, **Table S2**) in a 1 ml PCR reaction (300 μM dNTPs, 0.8 μM TSSR2, 0.8 μM TSSR-RPIX, 10 U Phusion polymerase in 1X Phusion HF buffer) for 2 cycles. The added 71bp allowed separation of the sequencing library from the adapter-less library on a 16% 0.5X TBE native polyacrylamide gel (19:1 Acrylamide/Bis-acrylamide) run at 200V. The 141bp band was excised, and the DNA recovered by electroelution as described above. The purity and concentration of the library was determined by Bioanalyzer (High-sensitivity dsDNA chip, Agilent). Multiple sequencing libraries with compatible barcodes were pooled in equimolar concentrations and sequenced on a HiSeq2000 with 10 million reads per library, using single-end, 50bp sequencing mode.

### Motif discovery

#### Raw sequencing data processing

The sequenced libraries were processed using the *R* package *SELEX* (http://bioconductor.org/packages/SELEX). We required sequencing reads to match the sequence TGGAA at positions 24-28. The package was also used to construct Markov models of R0 and compute the information gain (KL divergence) after affinity-based selection.

#### Model-based analysis (SelexGLM)

To avoid bias in our estimates, we split the reads into two equal-sized random subsets. One half was used to define the “universe” of unique variable regions (which we refer to as “probes”). Read counts across this universe were defined based on the other half of the reads, and a count of zero was registered for probes that were only seen in the first half. A Markov model of order 5 was constructed from the R0 probes using the selex.mm () function from the *SELEX* package, and an affinity table for k=15 was constructed using selex.affinities (). An initial position specific affinity matrix (PSAM) was constructed from the relative affinity of all 15x3 single-base mutations of the optimal 15-mer, expanded to the desired size by adding eight neutral columns on each side, and used as a seed. The subsequent iterative procedure alternated between two steps. First, the current PSAM was used to find the position/direction of highest affinity on either strand, the optimal “view” on the probe. If that optimal affinity was larger than 95% of the sum over all positions (including the top position), the probe was used in the analysis; otherwise, it was ignored. The set of optimal positions in each of the accepted probes was used to define a design matrix containing the base identity at each position relative to the start of the optimal view. Using the probe counts as independent variables, the logarithm of the expected probe frequency in R0 according to the Markov model as offset, and a logarithmic link function, a fit was performed using the glm () function. The regression coefficients were interpreted as free-energy differences ddG. All our analyses were implemented in the form of an *R* software package, which will be submitted to *Bioconductor* as soon as the manuscript has been accepted for publication. All computational figure panels in this paper were produced fully automatically from the raw sequencing data using *R* scripts that use the *SELEX* and *SelexGLM* packages.

### Isothermal Titration Calorimetry (ITC)

ITC was performed using a Microcal VP-ITC (GE) at 25°C. Protein and DNA samples were dialyzed into binding buffer (20 mM HEPES-KOH, pH 7.7, 250 mM KCl and 0.5 mM TCEP) at 4°C for 36∼48 hours before use. DNA samples were loaded into the syringe and titrated into the protein in the reaction cell. Each ITC experiment consisted of an initial 2μl injection, followed by 20 x 14.3 μl injections, with 240 seconds between injections. For AR-DBD, 50 μM DNA and 10 μM protein sample were used. For GR-DBD, 100μM DNA and 20 μM protein sample were used to increase signal. The raw isotherm was analyzed using NITPIC (Brautigam et al. 2016), followed by fitting to a one-site binding model in SEDPHAT (Brautigam et al. 2016). The mean and standard deviation of thermodynamic parameters were calculated based on at least three experimental replicates.

### ChIP-seq data analysis

Raw reads for AR (GSM759657 and GSM759658), GR (GSM759669), and IgG control (GSM759671) data from ChIP-seq experiments using LNCaP-1F5 cells were downloaded from Gene Expression Omnibus and aligned to the hg19 assembly using bowtie2 with settings ‘--sensitive --score-min L,-1.5,-0.3’. Peaks were called using macs2 width default settings and the “fold enrichment” (column 7) was used to quantify peak strength. The narrowPeak bed-file was filtered retain one entry per unique peak interval and the intervals were standardized to cover 200bp. The PSAM score were then calculated for each offset orientation of the peak and the largest value was recorded.

### Statistical analysis

Figure 1: To minimize the influence of sequencing count on the calculation of relative enrichment, we only used 15-mers with ≥100 counts in both R7 and R8 libraries. We consider the sequencing count of each k-mer as a random variable of poisson distribution, where the k-mer count (n) is the best estimation of the mean (λ). Therefore, the variance is approximated by the k-mer count, with an absolute error of Sqrt (n). A sequencing count ≥100 therefore restricts the absolute error no more than 10% of the mean (λ).

Figure 2C, 3B, 4, Figure S4, S5: To compare DNA binding affinities of GR and AR for different sequence by EMSA or ITC, we performed at least 3 independent experiments under the same conditions. The variance of measured affinity is assumed to be normally distributed, and the T-test and P-values are appropriate.

## DATA ACCESS

**The** raw SELEX-seq reads from this study have been submitted to the Sequence Read Archive (SRA; www.ncbi.nlm.nih.gov/sra) (Leinonen et al. 2011) under accession number SRP101815.

## ACKNOWLEDGMENTS

This research was supported by the National Institutes of Health (NIH) K99/R00 CA149088, the Roy J. Carver Charitable Trust 01-224, and the National Science Foundation CAREER grant 1552862 (MAP), and grant R01HG003008 (HJB). We thank Todd Riley and members of the Pufall, Bussemaker, Mann, and Rohs labs for useful discussions.

## AUTHOR CONTRIBUTIONS

M.A.P and H.J.B conceived and oversaw the project. L.Z designed and performed all experiments. G.D.M, J.F.K, C.R. and H.J.B conceived and developed the GLM analysis approach. G.D.M implemented the *SelexGLM* software and analyzed SELEX-seq data. H.T.R analyzed ChIP-seq data. L.Z and V.D.F performed preliminary analysis of SELEX-seq and ChIP-seq data. M.A.P analyzed the crystallography data. J.C.H provided the ITC hardware and consultation on interpretation. L.Z, G.D.M, H.T.R, H.J.B and M.A.P wrote the manuscript.

